# MetaFold-RNA: Accurate prediction of RNA secondary structure using a meta-learning-guided deep network

**DOI:** 10.1101/2025.09.18.676970

**Authors:** Lei Wang, Han Zhang, Xudong Li, Yan Wang, Zhidong Xue

**Author notes:** Correspondence author (Y.W.), (Z.X.). these authors contributed equally to this work (Lei Wang, Han Zhang).

## Abstract

Accurately predicting the secondary structure of ribonucleic acid (RNA) is a critical step toward deciphering its biological roles and engineering novel RNA-based technologies. However, achieving high accuracy and generalization, especially for unseen RNA families, has long remained a central challenge in computational biology. Here we present MetaFold-RNA, a novel meta-learning framework that sets a new state of the art in RNA secondary structure prediction. Instead of relying on a single predictive paradigm, MetaFold-RNA synergizes insights from disparate models through a meta-learning approach and refines predictions via a co-evolutionary optimization process. On the challenging bpRNA-new benchmark, which specifically tests performance on unseen RNA families, MetaFold-RNA surpasses all SOTA methods, outperforming the best previous approach by 7.3%. This breakthrough performance is consistently maintained across other stringent assessments, including the PDB-new dataset and blind CASP predictions. Furthermore, our method demonstrates high robustness by accurately resolving the complex topologies of artificial RNA nanostructures. This work establishes a new performance benchmark, particularly in cross-family generalization, and provides a powerful computational tool for accelerating the study and design of RNA.

## 1 Introduction

Ribonucleic acid (RNA) is a biopolymer of exceptional functional diversity, acting not only as a transient carrier of genetic information but also as a structural and catalytic scaffold central to processes like protein synthesis, gene regulation, and enzymatic reactions^1,2^. The function of an RNA molecule depends on the specific spatial structure it adopts upon folding. In this folding process, the RNA secondary structure—a map of base pairs formed by hydrogen bonds—serves as the foundational framework and plays a decisive role. It not only stabilizes the molecule’s core conformation but also directly constitutes many functional domains, a concept known as hierarchical folding^3^. Consequently, the accurate prediction of RNA secondary structure is a critical and often sufficient step for understanding RNA function and engineering novel RNA-based tools.

Experimentally, RNA secondary structures can be inferred at a large scale using chemo-enzymatic probing techniques coupled with high-throughput sequencing (e.g., SHAPE-seq)^4–7^. However, these methods provide constraints that are often sparse and noisy, offering indirect evidence of base pairing rather than a complete structure map. They typically lack the resolution to consistently and accurately determine the status of every nucleotide, especially within complex motifs or for nucleotides not amenable to modification. Consequently, a significant gap remains between the vast number of known RNA sequences and the limited availability of high-quality, experimentally determined secondary structures, which has spurred the development of computational prediction methods.

Currently, computational methods for RNA secondary structure prediction can be broadly categorized into three types: thermodynamic model-based methods, shallow machine learning methods, and deep learning methods. These categories clearly illustrate the field’s paradigm shift from physics-and-chemistry-based rules to data-driven learning. The first category, thermodynamic model-based methods, operates on the core assumption that an RNA molecule folds into its most thermodynamically stable conformation, which corresponds to the minimum Gibbs free energy. In these predictions, nested base-pairing structures are typically identified using dynamic programming strategies. Representative tools in this category include MFold/UNAfold^8,9^, ViennaRNA RNAfold^10^, and RNAstructure^11^. These methods rely on experimentally determined nearest-neighbor energy parameter sets, such as the Turner parameters^12–14^, to score and select optimal secondary structure conformations. While effective, they face several intrinsic limitations: first, their accuracy is bound by the completeness of the semi-empirical energy models. Second, computational complexity remains a bottleneck for very long sequences. Third, the underlying assumption of thermal equilibrium often neglects the complex, co-transcriptional folding dynamics. Finally, most dynamic programming frameworks are inherently designed for nested structures and struggle to predict non-nested interactions like pseudoknots.

The second category, shallow machine learning methods, emerged to overcome the limitations of purely thermodynamic models. These approaches no longer rely exclusively on fixed experimental energy parameters but instead employ statistical learning models to ‘learn’ a set of scoring parameters from known structural data. For instance, CONTRAfold^15^ and ContextFold^16^ predict structures based on sequence features. Another data-driven approach is EternaFold^17^, which uses a massive dataset from a citizen science platform to retrain and calibrate thermodynamic parameters. These methods paved the way for data-driven structure prediction but are relatively limited in their model capacity and automatic feature extraction capabilities. The third and most rapidly advancing category in recent years is deep learning (DL). For instance, SPOT-RNA^18^ utilizes a dual-channel architecture of convolutional neural networks (CNNs) and bidirectional long short-term memory networks (BiLSTMs). E2Efold^19^ pioneered an end-to-end approach. UFold^20^ innovatively encodes sequences as 2D images, applying U-Net models to extract multi-scale interaction information. More recent methods like NeuralFold^21^ and RFold^**?**^ have also introduced more advanced architectures. Furthermore, another class of methods is dedicated to integrating physical priors into the deep learning framework. For example, BPfold^22^ innovatively constructed a base-pair motif energy library, while MXfold2^23,24^ learns a scoring function fused with traditional energy parameters via a neural network.

Despite their impressive performance, deep learning models for RNA structure prediction remain highly sensitive to the distribution of training data, often exhibiting reduced robustness and limited generalization when applied to out-of-distribution (OOD) RNA families. In such cases, their predictive accuracy can degrade substantially, whereas thermodynamic models often maintain more stable performance. This brittleness reflects a fundamental limitation: overfitting to observed data and a failure to fully capture the universal biophysical principles governing RNA folding. To address this generalization gap, several state-of-the-art models have incorporated auxiliary information, typically through ad-hoc integrations. For instance, UFold leverages prior representations from CDPfold and predictions from CONTRAfold, including through knowledge distillation, to enrich its training data^20^. However, even with this distillation approach, UFold still exhibits lower accuracy than CONTRAfold in predicting RNA secondary structures across novel families. BPfold introduces a pre-computed library of base-pair motif energies as a physical prior^22^, and MXfold2 regularizes its predictions using a thermodynamically-informed loss function^24^. While these strategies offer incremental improvements, they do not fundamentally resolve the challenge of generalizing to novel RNA families. Consequently, developing models capable of robustly predicting structures for previously unseen RNA families remains a major unresolved bottleneck in the field.

To address this core generalization challenge, we present MetaFold-RNA, an architecture designed to synergistically fuse a meta-learning network with a powerful architectural inductive bias. At its foundation, our approach employs a meta-learning strategy by assimilating structural priors from a diverse ensemble of orthogonal prediction tools—including thermodynamic (LinearFold^25^), probabilistic (CONTRAfold^15^), and deep learning-based (UFold^20^, SPOT-RNA^18^) predictors. To further enrich these initial features and address the inherent sparsity of the pairing matrix, we also draw inspiration from CDPFold^26^ by incorporating an additional channel that reflects implicit base-pairing matches. This comprehensive initial representation is then processed by the core of MetaRNA-Fold model, which is built upon a stack of MetaFormer blocks. This architecture possesses a strong inductive bias specifically for co-evolving 1D sequence and 2D pair representations. This bias is realized through a dual-track system where information is iteratively exchanged: sequence features update the pairwise map via an outer product operation, and this map is in turn refined by specialized 2D convolutional modules. Critically, the refined structural map then informs the sequence representation through a pairwise-biased attention mechanism, directly guiding the model’s focus based on emerging structural patterns. Thus, MetaFold-RNA integrates the distilled knowledge from prior models (the meta-learning component) with a novel network structure designed to process it (the inductive bias), creating a powerful framework for accurate structure prediction.

We further demonstrate that MetaFold-RNA sets a new state of the art in predictive accuracy and, most critically, in generalization. In comprehensive benchmarks against leading methods on diverse datasets—including the stringent bpRNA-new cross-family test set, PDB-new, and recent CASP(15 and 16) challenges—MetaFold-RNA consistently achieves superior performance. Its robust generalization is further underscored by its remarkable success in accurately resolving the complex topologies of artificial RNA nanostructures, a task that tests a model’s grasp of biophysical principles beyond evolutionary data. This breakthrough in cross-family structure prediction not only establishes a new performance standard for the field but also provides a powerful computational tool to accelerate the exploration of the RNA structurome and the engineering of novel RNA-based technologies. To facilitate broad access to our method, we have developed the MetaFold-RNA web server, offering an intuitive interface for structure prediction and visualization.

## 2 Results

### 2.1 Overview of the MetaFold-RNA architecture

The MetaFold-RNA architecture is a deep neural network engineered for RNA secondary structure prediction. The model’s design is predicated on a dual-track information processing pipeline that explicitly disentangles and co-evolves a one-dimensional (1D) representation of the primary sequence and a two-dimensional (2D) representation of the base-pairing map (Fig. 1). This architecture facilitates iterative, bidirectional information flow between the two representations, enabling the synergistic modeling of sequential dependencies along the RNA backbone and the geometric constraints of the final folded structure. The computational workflow commences within an input module that generates these dual representations from the primary RNA sequence (Fig. 1a, b). The 1D sequence is projected into a high-dimensional feature space via a trainable embedding layer to yield the initial sequence representation (`seq_repr`). Concurrently, the initial pair representation (`pair_repr`) is constructed by integrating base-pairing probabilities derived from an ensemble of four distinct prediction algorithms, thereby providing a rich, multi-faceted structural prior. This 2D map is subsequently augmented with relative positional encodings to explicitly embed spatial information.

**Figure 1.**
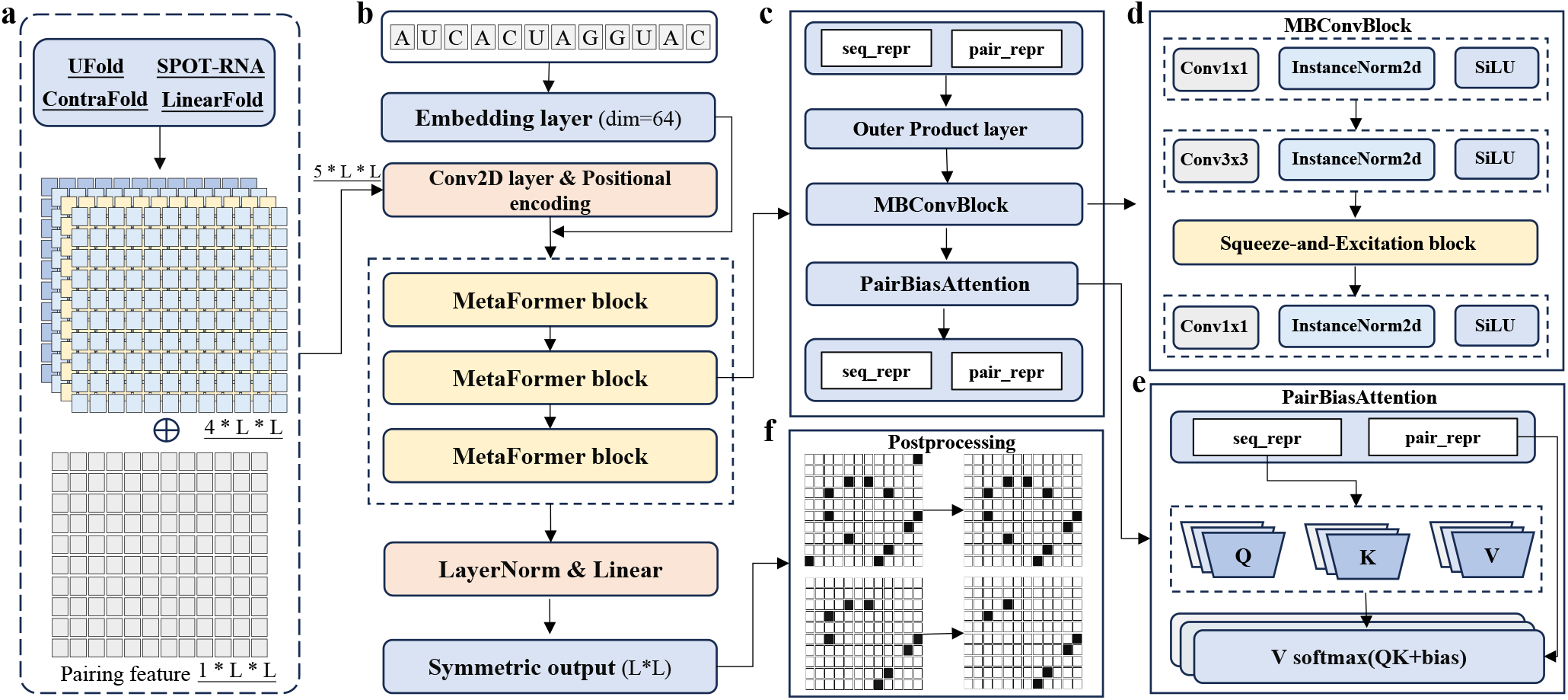
Overview of the MetaFold-RNA architecture. The model utilizes a dual-track system to process 1D sequence and 2D pairing representations. **a**, Initial inputs are generated from the RNA sequence (1D) and prior predictions from external tools (2D). **b**, The sequence is embedded, and this representation is processed to interact with the pair representation, which is augmented with positional encodings. **c**, A series of MetaFormer blocks iteratively exchange information between the sequence representation (`seq_repr`) and the pair representation (`pair_repr`). **d**, The MBConvBlock refines the 2D pair representation using efficient convolutional layers. **e**, The Pairwise-Biased Attention mechanism uses the refined 2D map to bias attention scores in the 1D sequence representation. **f**, The final pair representation is projected to a symmetric contact map and refined via post-processing to produce the final structure.

The central computational engine of MetaFold-RNA consists of a stack of three identical MetaFormer blocks that execute the co-evolutionary logic (Fig. 1c). Each block orchestrates a two-step information exchange. First, sequence-level information is propagated to the 2D map: an outer product operation transforms the 1D sequence embeddings into a pairwise representation, which is then fused with the existing `pair_repr`. This intermediate 2D representation is subsequently processed by a Mobile Inverted Bottleneck Convolution block (MBConvBlock, Fig. 1d), a computationally efficient module designed to refine spatial features. In the second step, this refined structural information is propagated back to the 1D sequence via a Pairwise-Biased Attention module (Fig. 1e). Here, the `pair_repr` directly provides a bias term to the attention scores, conditioning the self-attention calculation on the global structural context and enabling the model to dynamically allocate attention to nucleotide interactions supported by the evolving 2D map. Upon completion of the final block, the refined `pair_repr` is passed to a linear output head, generating a logit matrix representing the pairing likelihood for all residue pairs. This matrix is explicitly symmetrized to enforce the reciprocal nature of base pairing before a final, deterministic post-processing algorithm filters the predictions (Fig. 1f), removing any pairs that violate fundamental biophysical constraints to yield a physically valid secondary structure map.

### 2.2 Superior Performance on Within-Family Benchmarks

To conduct a rigorous evaluation of our proposed method, MetaFold-RNA, we performed a benchmark analysis against 14 contemporary methods on the independent test set TS0. These competing methods are broadly classifiable into three paradigms: deep learning models, thermodynamic models, and shallow machine learning models. The performance was assessed using three standard metrics: precision, recall, and the F1-score.

As illustrated in Figure 2a, MetaFold-RNA achieves an F1-score of 0.725, establishing a new state-of-the-art in prediction accuracy. Within the deep learning paradigm, which exhibits a wide performance spectrum, MetaFold-RNA emerges as a distinct top-performer. It surpasses the next best-performing methods, BPfold (F1-score = 0.658) and UFold (F1-score = 0.654), by a substantial margin of 10.2% and 10.8%, respectively. The foundation of this superior performance is an exceptional balance between precision and recall. MetaFold-RNA attains the highest recall (0.815) across all tested methods, indicating a superior sensitivity for identifying true native base pairs, which is critical for functional analysis. Concurrently, it maintains a highly competitive precision of 0.676, ensuring that the predictions are not overwhelmed by false positives. This profile contrasts sharply with other top-tier deep learning models that exhibit a more pronounced trade-off; for instance, RFold achieves the highest precision (0.692) but at the expense of a significantly lower recall (0.635). Moreover, the performance of MetaFold-RNA constitutes a paradigm shift compared to traditional methods. The F1-scores of thermodynamic models (e.g., RNAStructure at 0.533, LinearFold at 0.550) and shallow learning models (e.g., ContextFold at 0.546, ContraFold at 0.567) are largely confined below the 0.6 threshold. The commanding lead of MetaFold-RNA thus underscores the power of its architecture to decipher complex structural information that remains elusive to previous generations of algorithms.

**Figure 2.**
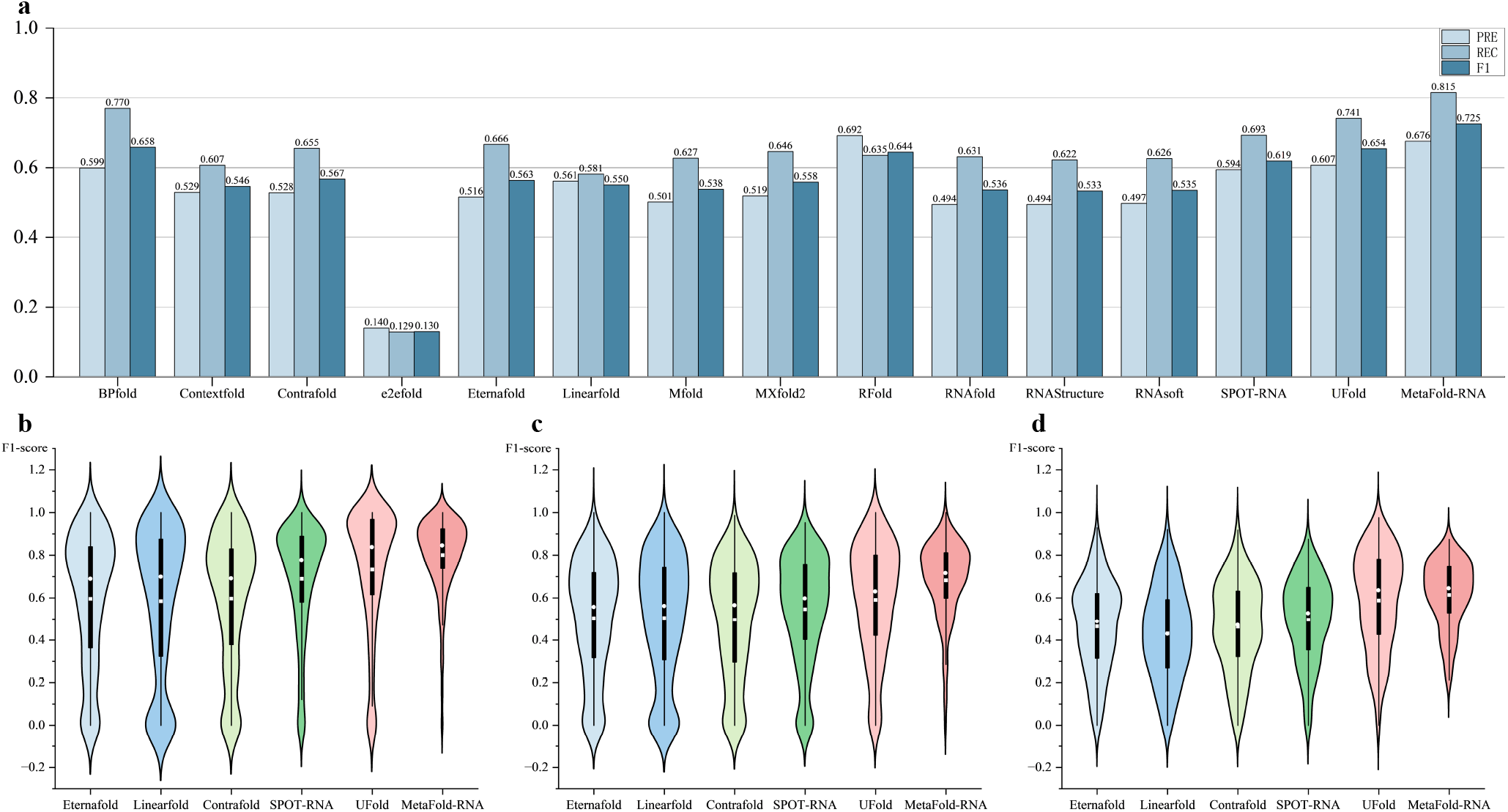
Performance comparison of MetaFold-RNA with other methods on the TS0 dataset. **a**, Bar chart showing overall precision, recall, and F1-scores. **b, c**, and **d**, Violin plots comparing F1-score distributions for top-performing methods on sequences of length 1-100 nt, 101-200 nt, and 201+ nt, respectively.

To further probe the robustness and reliability of our method, we analyzed the full distribution of F1-scores on sequence subsets of varying lengths. For this, we selected a representative cohort of methods from Figure 2a, covering the different paradigms. The violin plots in Figure 2b-d visualize these distributions. Beyond simple central tendencies, the shape of the violins reveals crucial information about prediction consistency. For short sequences (1-100 nt, Figure 2b), the distribution for MetaFold-RNA is noticeably more compact and skewed towards higher F1-scores, with a denser concentration of results near its high median of 0.845. This indicates a higher consistency in producing accurate predictions compared to other methods, whose distributions are wider and more symmetric. As sequence length increases to 101-200 nt (Figure 2c), the performance of all methods degrades, yet the distribution for MetaFold-RNA remains more favorably skewed than its competitors. This robustness is most striking for long sequences (201+ nt, Figure 2d), a known bottleneck for secondary structure prediction. Here, the distributions for many methods exhibit long lower tails, signifying a higher frequency of poor-quality predictions. In contrast, the distribution for MetaFold-RNA is more contained, with fewer extreme outliers at the low end. This demonstrates not only a higher average performance (mean F1-score of 0.614) but, more importantly, a greater reliability in handling complex, long-range dependencies. To evaluate the ability of MetaFold-RNA to capture global structural dependencies, we further assessed its performance on long-range base pairs, defined as those separated by more than half the sequence length (*L/*2). As shown in Supplementary Table 1, MetaFold-RNA achieves the highest F1-score (0.741) among all methods, with a precision of 0.729 and recall of 0.858. These results indicate that MetaFold-RNA accurately identifies distant base-pairing interactions with high sensitivity and precision. In comparison, UFold and SPOT-RNA achieve F1-scores of 0.676 and 0.634, respectively, while traditional models such as LinearFold (0.545) and ContraFold (0.558) perform substantially worse. These findings highlight the advantage of MetaFold-RNA in modeling non-local interactions, which are essential for capturing complex RNA secondary structures. Having established its state-of-the-art performance on familiar data, the crucial next test is to assess whether this capability generalizes to unseen RNA families.

**Table 1.** Performance on long-range base pair prediction (sequence separation > *L/*2). This table shows a representative subset of high-performing models. The best F1-score in each dataset is highlighted in bold.

| Method | CASP15 |  |  | CASP16 |  |  | PDB_new |  |  |
| --- | --- | --- | --- | --- | --- | --- | --- | --- | --- |
|  | Precision | Recall | F1 | Precision | Recall | F1 | Precision | Recall | F1 |
| LinearFold | 0.782 | 0.886 | 0.823 | 0.960 | 0.615 | 0.594 | 0.806 | 0.610 | 0.546 |
| ContraFold | 0.773 | 0.909 | 0.829 | 0.739 | 0.830 | **0.703** | 0.675 | 0.803 | 0.686 |
| SPOT-RNA | 0.809 | 0.706 | 0.658 | 0.706 | 0.776 | 0.640 | 0.729 | 0.771 | 0.679 |
| UFold | 0.807 | 0.751 | 0.747 | 0.597 | 0.698 | 0.600 | 0.622 | 0.709 | 0.597 |
| EternaFold | 0.776 | 0.909 | 0.830 | 0.629 | 0.869 | 0.656 | 0.612 | 0.815 | 0.657 |
| **MetaFold-RNA** | 0.902 | 0.879 | **0.870** | 0.741 | 0.796 | 0.671 | 0.790 | 0.816 | **0.734** |

To quantify the benefit of our meta-learning approach, we benchmarked MetaFold-RNA directly against the four models it leverages as priors: LinearFold, Contrafold, SPOT-RNA and UFold (Figure 3). MetaFold-RNA achieves a higher F1 score than Contrafold in 81.6% of test cases, SPOT-RNA in 81.5%, LinearFold in 75.2%, and UFold in 57.7%. These results confirm that MetaFold-RNA not only aggregates prior knowledge effectively, but also generalizes well across modeling assumptions. The strongest performance margins—over Contrafold and SPOT-RNA—suggest that the model benefits most from combining complementary signals from probabilistic and deep learning predictors. The more modest gain over UFold, another data-driven deep-learning model, highlights the challenge of surpassing high-capacity architectures, but also underscores the residual benefits of structured fusion. Altogether, the findings highlight the integrative strength of MetaFold-RNA in reconciling heterogeneous predictive results into a unified and accurate structure predictor.

**Figure 3.**
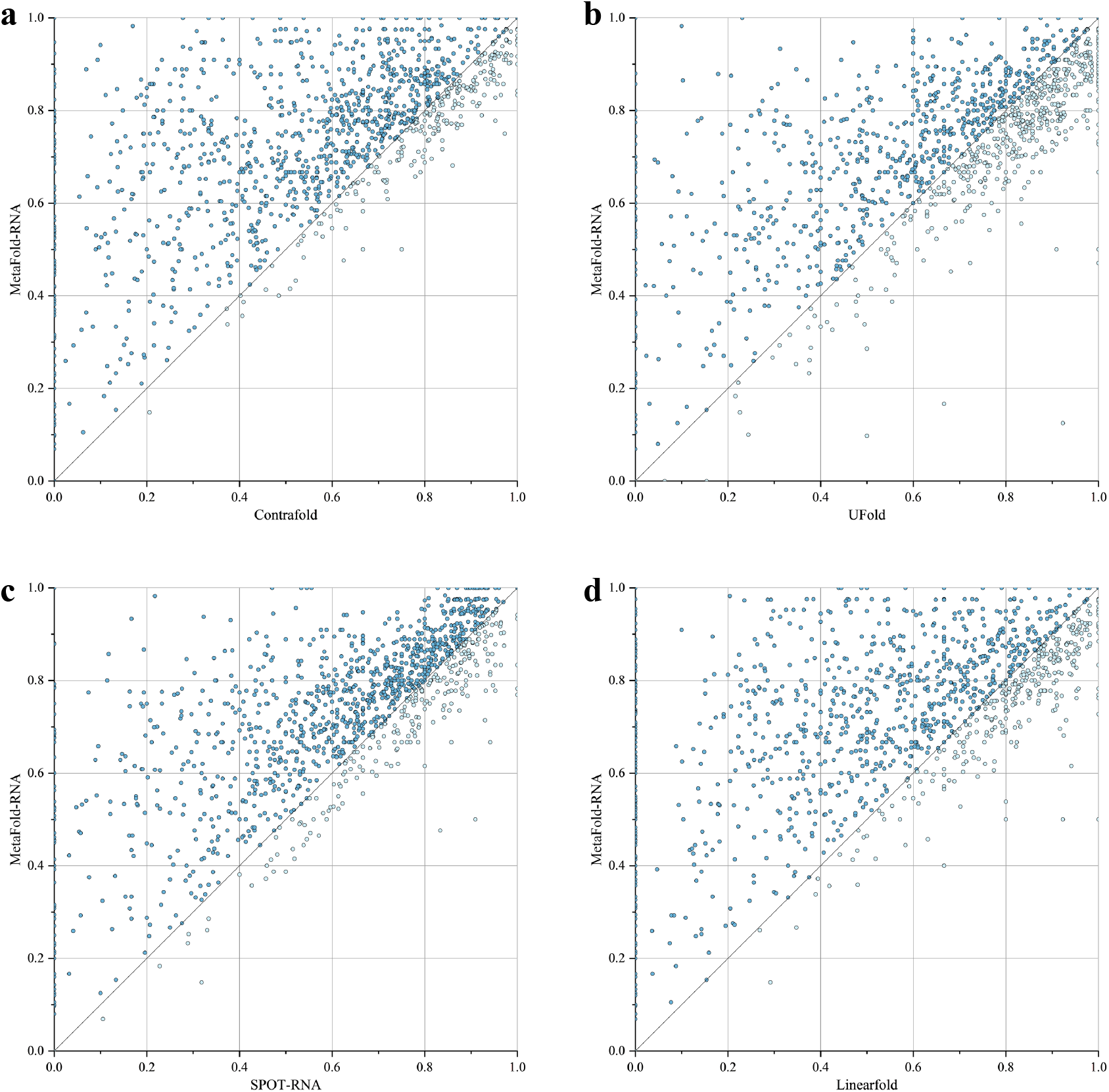
Head-to-head comparison between Metafold-RNA and baseline models on the TS0 dataset. Each panel shows the proportion of test samples in which Metafold-RNA outperforms a specific model in terms of secondary structure prediction accuracy. (a) Contrafold, (b) UFold, (c) SPOT-RNA, (d) LinearFold.

### 2.3 Robust Generalization Across Unseen RNA Families

Building on the strong baseline results from the TS0 set, we next subjected MetaFold-RNA to a more demanding test of generalization using the bpRNA-New dataset, which consists of RNA sequences from entirely unseen families during training. This dataset provides a stringent benchmark for assessing cross-family prediction performance. We compared MetaFold-RNA against 14 competing algorithms spanning three major paradigms: deep learning-based models (e.g., UFold, SPOT-RNA), thermodynamic models (e.g., RNAFold, RNAStructure), and shallow machine learning methods (e.g., ContraFold, ContextFold). Evaluation was conducted using standard metrics of precision, recall, and F1-score, as shown in Figure 4a.

**Figure 4.**
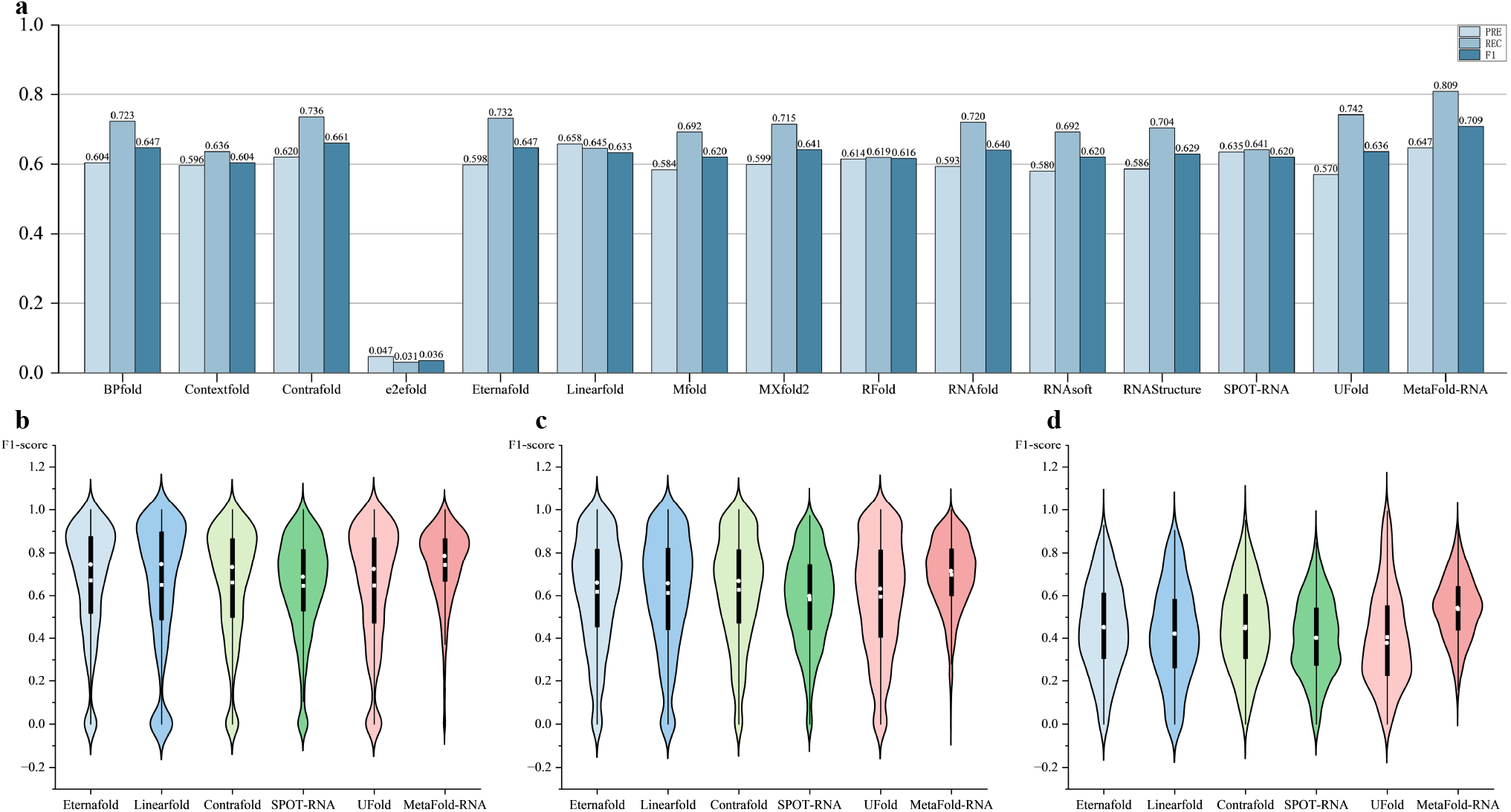
Cross-family prediction performance on the bpRNA-New dataset. (**a**) Overall precision, recall, and F1-score of 15 RNA secondary structure prediction methods evaluated on unseen RNA families. (**b–d**) Violin plots of F1-scores for representative methods stratified by sequence length: (**b**) 1–100 nt, (**c**) 101–200 nt, (**d**) 201+ nt.

MetaFold-RNA achieves the highest F1-score of 0.709, significantly outperforming all baselines. This superiority is driven by the highest recall (0.809), reflecting its remarkable sensitivity in identifying native base pairs in previously unseen RNA families, while maintaining a high precision of 0.647. The second-best method, ContraFold, reaches an F1-score of 0.661, followed closely by BPfold and EternaFold (both at 0.647). However, these models diverge in their balance: BPfold emphasizes recall (0.723) at the cost of precision, whereas EternaFold achieves slightly better precision (0.598) but with lower recall. Among deep learning models, UFold (F1 = 0.636) shows strong recall (0.742) but suffers from the lowest precision (0.570), leading to many false positives. SPOT-RNA (F1 = 0.620) and MXFold2 (F1 = 0.641) perform moderately, while E2EFold exhibits a dramatic drop (F1 = 0.036), indicating a failure to generalize across families. Compared to the TS0 dataset, most models experience a performance degradation on bpRNA-New, highlighting their limited generalization. In contrast, MetaFold-RNA maintains an optimal balance between precision and recall, suggesting it captures deeper structural principles that extend beyond family-specific features.

To further evaluate prediction robustness across structural complexity, we analyzed the distribution of F1-scores by stratifying sequences into three length-based groups: short (1–100 nt), medium (101–200 nt), and long (201+ nt). For each group, we visualized prediction performance using violin plots (Figure 4b–d), which capture both the central tendencies and the variability of model outputs. In the short-sequence group (Figure 4b), most methods achieve relatively high F1-scores, but their distributions differ notably. MetaFold-RNA exhibits a compact, sharply peaked distribution centered near its median (0.786), indicating not only strong but also consistent performance. By contrast, models such as UFold and SPOT-RNA show wider distributions, suggesting unstable performance even for shorter RNAs. In the medium-length group (Figure 4c), performance begins to diverge further. MetaFold-RNA retains a high median F1-score (0.713) and a narrow distribution, whereas other methods begin to show flattened violins and heavier tails, implying increased frequency of poor predictions. In the long-sequence regime (Figure 4d), structural complexity poses a major challenge. Here, MetaFold-RNA’s violin remains relatively contained, with a median F1-score of 0.542—substantially higher than competitors, whose violins stretch downward with long tails and low-density cores. UFold, SPOT-RNA, and EternaFold in particular show pronounced distributional spread, indicating frequent performance collapse on complex long RNAs. Together, these distributional trends reinforce the conclusion that MetaFold-RNA not only outperforms competing methods in average and median metrics, but also delivers more stable predictions across diverse sequence lengths.

To further probe MetaFold-RNA’s ability to model global structural information under distributional shift, we evaluated its performance on long-range base pairs, defined as those separated by more than half the sequence length (*L/*2). As shown in Supplementary Table 2, MetaFold-RNA again achieves the highest F1-score (0.753). This result confirms that even on unfamiliar RNA families, MetaFold-RNA is exceptionally capable of identifying distant base-pairing interactions with high confidence and sensitivity. In comparison, other methods show a considerable performance gap. For instance, the next-best competitors, LinearFold and ContraFold, posted F1-scores of 0.677 and 0.663, respectively. Other deep learning models, such as UFold (0.613) and SPOT-RNA (0.598), lagged even further behind. These findings underscore that the architectural advantages of MetaFold-RNA in processing non-local dependencies are robust and generalize effectively, making it a highly reliable tool for deciphering the complex, global topologies of novel RNAs.

To precisely quantify the benefits of our meta-learning approach under distributional shift, we conducted a head-to-head comparison of MetaFold-RNA against the four models it uses as priors on the bpRNA-New dataset (as shown in Figure 5). The analysis reveals that MetaFold-RNA demonstrates a powerful ability to integrate and refine predictions, even when faced with RNA sequences from unseen families. Specifically, MetaFold-RNA achieves a higher F1-score than **SPOT-RNA** in **71.9%** of test cases, outperforms **UFold** in **63.8%**, surpasses **ContraFold** in **62.5%**, and exceeds **LinearFold** in **58.1%** of cases. These figures robustly demonstrate that MetaFold-RNA effectively synthesizes signals from disparate predictor types—including deep learning, shallow machine learning, and thermodynamic models—and distills them into a more accurate and generalizable prediction than any single model can offer. This result reaffirms that the superior performance of MetaFold-RNA stems not only from learning on the training data but also from an architecture that captures universal structural principles, allowing it to maintain its lead in the stringent challenge of cross-family prediction. This robust performance on both within- and cross-family data demonstrates the model’s powerful learning capacity. To complete our evaluation, we assessed its performance in the most realistic and challenging scenarios: blind prediction against recently determined experimental structures.

**Figure 5.**
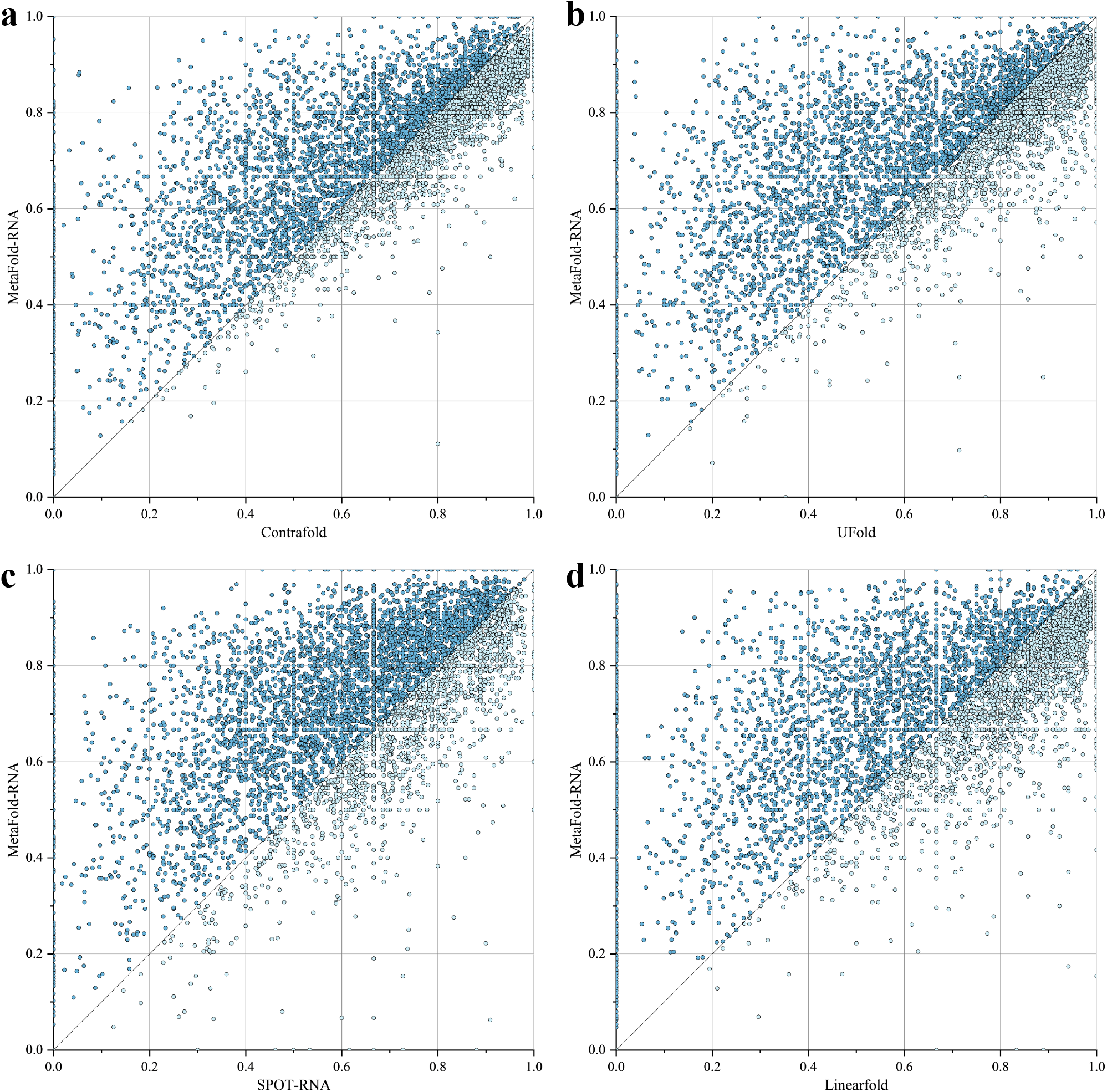
Head-to-head F1-score comparison on the bpRNA-New cross-family dataset. Each panel is a scatter plot comparing the performance of MetaFold-RNA against one of the four models used as priors. Points located above the diagonal line indicate a higher F1-score for MetaFold-RNA. The comparisons are against: (a) Contrafold, (b) UFold, (c) SPOT-RNA, and (d) LinearFold.

### 2.4. Performance Evaluation on CASP and PDB Benchmarks

In the final and most stringent stage of our evaluation, we measured MetaFold-RNA’s performance against the gold standard of blind prediction challenges: the CASP15 and CASP16 experiments, alongside a forward-looking test on the PDB-new dataset. Among these, CASP15 (n=11) and CASP16 (n=13) serve as the gold standard for testing predictive power on unknown structures, as they are recent, official blind assessment experiments. In the CASP15 test (Figure 6a), MetaFold-RNA achieved the highest F1-score of 0.818. Among the 14 competing tools, the five **traditional thermodynamic models** had F1-scores ranging from 0.682 to 0.816, with LinearFold (0.816) being highly competitive and nearly matching MetaFold-RNA. The two **shallow learning models** scored robustly between 0.759 and 0.809. In contrast, the seven **deep learning competitors** exhibited immense performance variance, with F1-scores spanning from a low of 0.028 (e2efold) to a high of 0.814 (MXfold2), revealing the paradigm’s great potential and instability. In CASP16 (Figure 6b), MetaFold-RNA’s lead (0.780) was more pronounced. Interestingly, the top-performing competitor was a shallow learning model, Contextfold (0.753); traditional models remained consistent (range 0.648–0.726); and the deep learning category again showed vast divergence (from 0.062 to 0.679). These blind tests collectively show that while MetaFold-RNA is consistently at the top, leading traditional and shallow learning methods remain formidable competitors.

**Figure 6.**
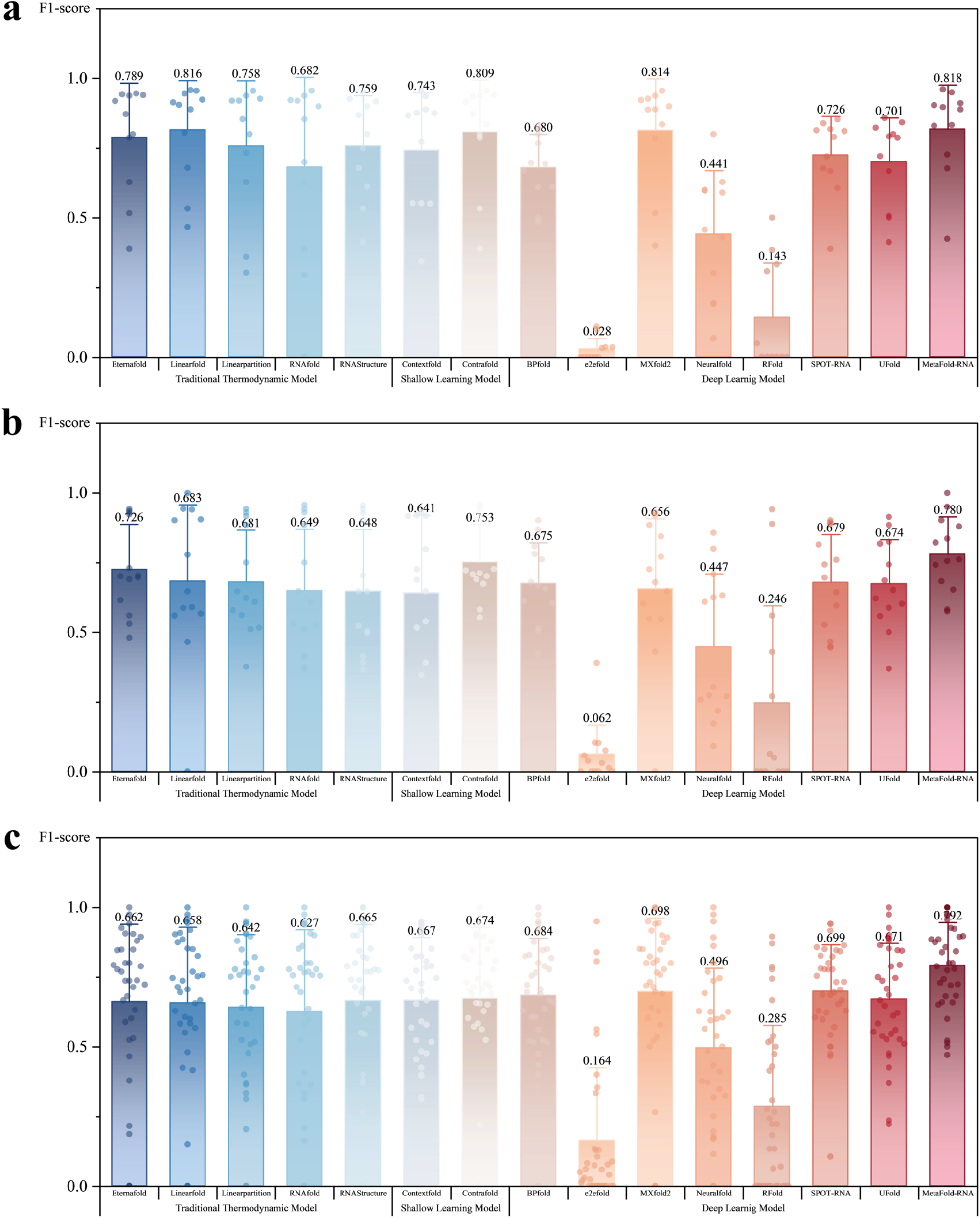
Comparison of MetaFold-RNA and competing methods in terms of F1-score across three independent datasets. Panel (**a**) shows results on CASP15, panel (**b**) on CASP16, and panel (**c**) on PDB-new. MetaFold-RNA consistently achieves the highest performance across all benchmarks.

To assess robustness against future data, we used the PDB-new dataset (n=36), which contains the latest PDB structures as of 2025 and is key for measuring generalization. In this test (Figure 6c), MetaFold-RNA’s superiority was most evident, with an F1-score (0.792) that was over 13% higher than any other tool. The performance distribution of paradigms shifted significantly: traditional thermodynamic (F1 range: 0.627–0.662) and shallow learning (F1 range: 0.665–0.674) models were grouped into a mid-tier performance bracket. In stark contrast, deep learning models were absolutely dominant, with the top three performers—MetaFold-RNA, SPOT-RNA (0.699), and MXFold2 (0.698)—all belonging to this category. This clearly demonstrates the paradigm’s advantage in learning and predicting novel, complex structures. To further probe the ability to capture global architecture, we compared representative models on long-range base-pair prediction (Table 1). MetaFold-RNA ranked first on both CASP15 and PDB-new, confirming its strong capability. Although ContraFold performed slightly better on CASP16, this highlights the subtle advantages of different architectures on specific tasks. In conclusion, while traditional and shallow learning methods are reliable tools, the state-of-the-art deep learning approach represented by MetaFold-RNA shows superior robustness and generalization on the newest, most complex structures, particularly in resolving the long-range dependencies that define global architecture, marking a clear path forward for the field.

### 2.5. Visual Analysis of Prediction Performance on Synthetic CASP15 Targets

A primary challenge for existing RNA secondary structure prediction methods, which are predominantly trained and bench-marked on natural RNAs, is their ability to generalize to synthetic and engineered constructs. These artificial structures often feature non-canonical motifs and designed global architectures not prevalent in the training data, thus serving as a rigorous test of whether a model has learned the fundamental principles of RNA folding, rather than merely recognizing patterns. To demonstrate MetaFold-RNA’s capabilities in this demanding area, we performed a comparative visual analysis on two representative synthetic RNAs from the CASP15 experiment (Figure 7).

**Figure 7.**
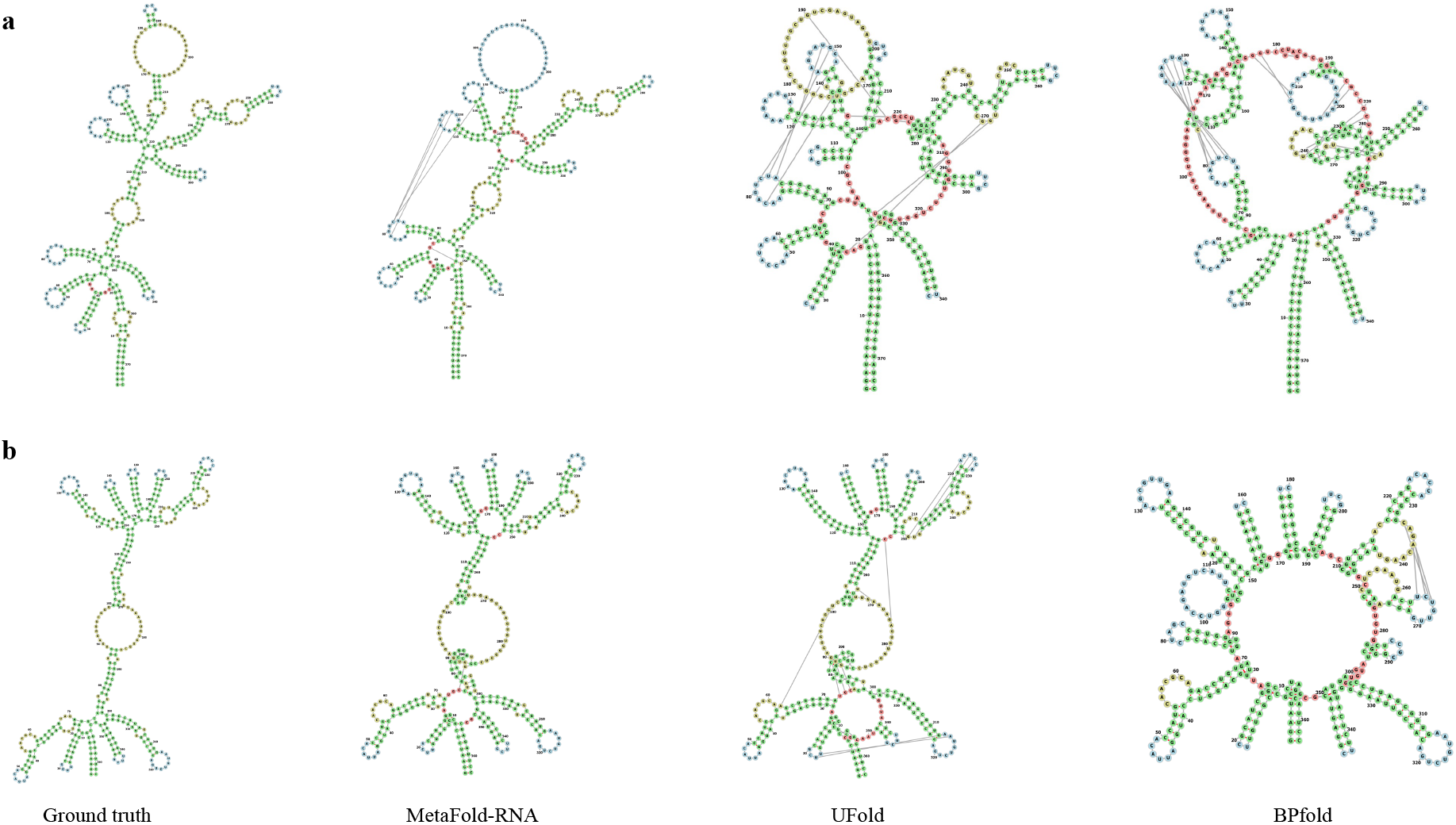
Visualization of secondary structure predictions on challenging CASP15 targets. (**a**) The Ligand-bound state of a broccoli-pepper aptamer FRET tile (R1136). (**b**) A synthetic RNA origami nanodevice, the Traptamer (R1126).

The first case is R1136 (Figure 7a), a Ligand-bound state of a broccoli-pepper aptamer FRET tile. On this target, MetaFold- RNA achieved a high F1-score of 0.911, significantly outperforming two other state-of-the-art deep learning methods, UFold (0.721) and BPfold (0.669). The structural visualization clearly corroborates this quantitative difference: the prediction by MetaFold-RNA closely recapitulates the ground truth topology. The second case is R1126 (Figure 7b), an RNA origami 3-helix tile nanorobot engineered as a “Traptamer”. On this highly challenging target, MetaFold-RNA achieved a near-perfect F1-score of 0.950, again substantially leading BPfold (0.694) and showing a clear advantage over the strong competitor UFold (0.858). The visualization (Figure 7b) intuitively shows that MetaFold-RNA precisely reconstructed the multi-helix topological structure of this RNA nanodevice. These two cases, particularly the successful prediction of highly designed synthetic RNAs, provide strong evidence of MetaFold-RNA’s superior ability to decipher complex structural patterns compared to other leading methods.

## 3. Discussion

In this work, we introduced MetaFold-RNA, a novel meta-learning framework that sets a new state-of-the-art for RNA secondary structure prediction. Our comprehensive benchmarking across a diverse suite of datasets, including established benchmarks (TS0, bpRNA-New) and rigorous blind tests (CASP15, CASP16, PDB-new), demonstrates that MetaFold-RNA consistently and substantially outperforms existing methods from all major paradigms—thermodynamic, shallow machine learning, and deep learning. The key to this success lies in a synergistic combination of two core innovations: a meta-learning strategy for integrating diverse structural priors and a co-evolutionary architecture with a strong inductive bias for modeling the interplay between sequence and structure.

A central pillar of MetaFold-RNA’s design is the principle of knowledge integration through meta-learning. Rather than relying on a single source of information or a single algorithmic philosophy, our model assimilates initial structural hypotheses from an ensemble of orthogonal predictors (LinearFold, CONTRAfold, UFold, and SPOT-RNA). This approach yields a rich and robust initial representation that mitigates the inherent biases and weaknesses of any individual method. For instance, thermodynamic models may falter on structures with kinetics-driven folds not at a free energy minimum, while some deep learning models may struggle with RNA families underrepresented in the training data. By fusing these heterogeneous signals, MetaFold-RNA starts its refinement process from a more informed and balanced baseline. Our head-to-head comparisons (Figure 3 and Figure 5) empirically validate this strategy, showing that MetaFold-RNA consistently improves upon the predictions of its constituent priors, demonstrating a true synergistic effect rather than simple aggregation.

The second, and arguably more critical, innovation is the co-evolutionary network architecture. Traditional deep learning pipelines often process sequence and structural information in a unidirectional or loosely coupled manner. In contrast, MetaFold-RNA employs a dual-track system that facilitates iterative, bidirectional communication between the 1D sequence representation and the 2D pair representation. This is achieved through two specialized mechanisms. First, the outer product operation explicitly projects learned 1D sequence features into the 2D pairwise space, directly updating the structural map with sequential context. Second, and more importantly, the Pairwise-Biased Attention mechanism reverses this flow of information. It uses the refined 2D structural map to directly bias the attention scores in the 1D sequence module. This creates a powerful feedback loop: an emerging structural hypothesis guides the model to focus on the most relevant nucleotide interactions, which in turn leads to a more refined structural map in the next iteration. This architectural inductive bias, which closely mirrors the physical reality that local sequence motifs guide global folding and the emerging global fold constrains local interactions, is a primary driver of our model’s high accuracy, particularly in resolving complex, long-range dependencies (Table 1).

The superior performance of MetaFold-RNA is most striking in its generalization ability. On the cross-family bpRNA-New dataset, where many methods trained on specific families suffer a performance drop, MetaFold-RNA maintains its high accuracy (Figure 4). This suggests that our model has learned more fundamental, generalizable principles of RNA folding rather than merely memorizing family-specific patterns. This robustness is further corroborated by its top-ranking performance on the most recent PDB structures (PDB-new) and the challenging blind CASP assessments. The visual analysis of synthetic RNA constructs from CASP15 (Figure 7), such as the `Traptamer` nanostructures, provides compelling evidence that MetaFold-RNA can successfully decipher the structure of highly engineered sequences with topologies not commonly seen in natural RNAs, a feat that challenges many existing algorithms.

Despite its strong performance, MetaFold-RNA has several limitations that point toward future avenues of research. First, its performance is still indirectly dependent on the quality of the initial predictions from the prior models. While it demonstrates an ability to correct and improve upon them, its ceiling may be constrained if all priors provide poor initial hypotheses. Future work could explore dynamic weighting schemes or alternative methods for generating initial structural seeds. Second, like most current high-performance methods, our model is primarily designed to predict canonical and wobble base pairs, and it does not handle non-nested interactions such as pseudoknots. Extending the architecture to predict these more complex topologies is a critical next step for the field. Finally, the current framework operates de novo. Integrating experimental data, such as SHAPE or DMS probing information, as an additional input channel could further constrain the conformational search space and enhance prediction accuracy, especially for structurally ambiguous regions.

In conclusion, MetaFold-RNA represents a significant advance in computational RNA biology. By uniquely combining a meta-learning strategy with a co-evolutionary architecture, it robustly and accurately predicts RNA secondary structure across a wide range of biological and synthetic contexts. Its superior performance, especially in generalization and handling long-range interactions, establishes a new benchmark for the field. Through the public web server, we hope MetaFold-RNA will serve as a valuable tool for the broader research community, accelerating the study of RNA structure-function relationships and aiding in the design of novel RNA-based therapeutics and nanotechnologies.

## 4. Methods

### 4.1. Datasets

To comprehensively train and evaluate our model, we constructed multiple datasets for base training, transfer learning, and multi-dimensional testing.

#### Training Dataset

We integrated two widely used RNA secondary structure datasets—RNAStralign^27^ and bpRNA-1m^28^—to build the large-scale training set, following the strategy of state-of-the-art models such as UFold and BPfold. Specifically, 20,923 non-redundant sequences were selected from RNAStralign, covering eight major RNA families, while 10,814 representative sequences were randomly sampled from bpRNA-1m after rigorous filtering. These were merged and deduplicated using cd-hit-est^29^ with an 80% sequence identity threshold, resulting in a final set of 12,628 unique sequences.

#### Transfer Learning Dataset

Our transfer learning strategy aligns with state-of-the-art methods such as SPOT-RNA^18^ and UFold^20^. The primary motivation for this approach, as highlighted in the SPOT-RNA paper, is that widely-used, large-scale RNA secondary structure databases, such as bpRNA, predominantly rely on automated annotation pipelines. Consequently, the accuracy of these annotations has inherent limitations when compared to secondary structures derived directly from experimentally resolved structures. To circumvent the potential noise and inaccuracies introduced by automated annotations and to provide our model with training data based on an experimental “gold standard,” we opted to fine-tune it using a more precise, smaller dataset. Therefore, we chose as our foundation the same set of 120 RNA sequences from the PDB that was also utilized in the SPOT-RNA study. We re-downloaded the structural data for these entries and extracted their secondary structures using the forgi tool^30^. To ensure the quality and integrity of the dataset, any sequence exceeding 1400 nucleotides or any structure with missing residues in its PDB file was excluded. This rigorous curation process yielded a final dataset of 114 high-quality sequences for transfer learning. This approach ensures that our model is exposed to highly reliable base-pairing information during the final fine-tuning stage.

#### Test Datasets

We adopted four independent test sets to evaluate different models:

- **TS0**: 1,305 sequences from bpRNA-1m, partitioned as in MXfold2^24^, to evaluate performance on familiar RNA families.
- **bpRNA-new**: 5,401 sequences from novel RNA families, as defined in^24,31,32^, to assess generalization capabilities.
- **CASP**: Comprising 11 released RNA targets from CASP15 and 13 from CASP16. These targets represent challenging prediction tasks and have not been filtered by length.
- **PDB-new**: A dataset of 36 RNA chains from PDB entries published after 2025, with a resolution better than 3.5 Å. Redundancy was removed using cd-hit-est at an 80% sequence identity cutoff.

The length distribution of these datasets is visualized in Supplementary Figure 1. This highlights the diversity of sequence lengths across the test sets, ranging from under 100 nt to over 200 nt, which ensures a comprehensive evaluation of the model’s performance on RNAs of various sizes.

### 4.2. The network of MetaFold-RNA

We propose a deep learning model named MetaFold-RNA for the accurate prediction of RNA secondary structure. The essence of this task is to map a one-dimensional (1D) base sequence to a two-dimensional (2D) base-pairing map. Therefore, the core design philosophy of our model is to explicitly separate and facilitate iterative interaction between the 1D sequence representation and the 2D pairing representation. By constructing a dual-track information processing pipeline, the model can simultaneously capture the contextual dependencies of the sequence and the spatial geometric relationships of the pairings, with specialized modules enabling efficient communication between the two tracks. Our model consists of three cascaded core components: a Co-evolution Block, which is flanked by an input module for data preprocessing and an output module for generating the final prediction.

#### 4.2.1. Input Feature Representation and Encoding

Given an RNA sequence of length *L*, denoted as **x** = [*x*_1_, *x*_2_, …, *x*_*L*_], where *x*_*i*_ ∈ {A, U, C, G}, the model first maps each nucleotide to a continuous vector through a learnable embedding layer, resulting in an initial sequence representation 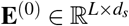, where *d*_*s*_ = 64.

To incorporate structural information, we integrate both handcrafted pairwise features (*C*_1_ = 1) and base-pairing probabilities predicted by four R A secondary structure prediction tools (*C*_2_ = 4). Specifically, UFold and SPOT-RNA are state-of-the-art deep learning-based methods, LinearFold uses thermodynamic energy minimization, and ContraFold is based on shallow probabilistic modeling. These predicted contact maps are concatenated and projected into a unified pairwise embedding space of dimension *d*_*p*_ = 32 via a 1×1 convolution:

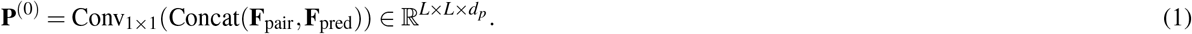

To further inject spatial context, we compute the relative distance Δ_*i j*_ = *i* − *j* for each base pair (*i, j*), discretize it into 65 bins over the range [−32, 32], and apply one-hot encoding:

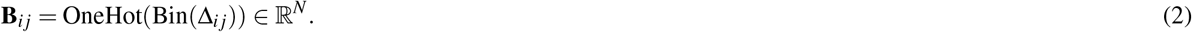

The positional encoding is then linearly projected using a learnable weight matrix 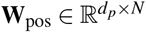, and added to the pairwise embedding:

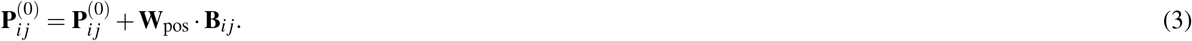

#### 4.2.2. MetaFormer: A Co-evolutionary Architecture for Sequence and Pair Representations

The core of our architecture is a stack of *N* = 3 co-evolutionary blocks, each combining 1D transformer-like processing with 2D CNN-based refinement and explicit modeling of co-evolution through outer product features. Each block *k* takes the representations **E**^(*k*)^ and **P**^(*k*)^ as input and produces refined representations **E**^(*k*+1)^ and **P**^(*k*+1)^.

##### (a) Sequence-to-pair information flow

Information from the sequence representation is used to update the pairwise representation. First, **E**^(*k*)^ is normalized via LayerNorm to produce 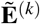. The outer product of the normalized feature vectors for each residue pair (*i, j*) is computed and subsequently projected to the pair dimension *d*_*p*_ to form an update tensor **P**^outer^:

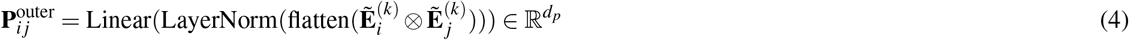

This update is then added to the input pair representation: **P**^′(*k*)^ = **P**^(*k*)^ + **P**^outer^.

##### (b) Pairwise representation refinement

The representation **P**^′(*k*)^ is refined using a Mobile Inverted Bottleneck Convolution (MBConv) block (Figure 1d). The tensor first undergoes LayerNorm and is then processed by the MBConv block, which consists of: (i) a 1×1 convolution for channel expansion, followed by InstanceNorm2d and a SiLU activation; (ii) a 3× 3 depthwise separable convolution, also followed by InstanceNorm2d and SiLU; (iii) a Squeeze-and-Excitation module for channel-wise feature recalibration; and (iv) a final 1 × 1 projection convolution with InstanceNorm2d. The output is passed through a dropout layer and added back to **P**^′(*k*)^ via a residual connection to yield the block’s final pair representation:

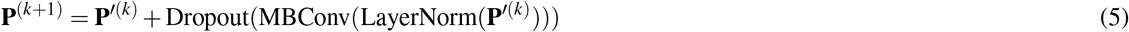

##### (c) Updating Sequence Representation with Pairwise-Biased Attention

The refined pair representation **P**^(*k*+1)^ is used to update the sequence representation via a Pairwise-Biased Attention mechanism (Figure 1e). Both **E**^(*k*)^ and **P**^(*k*+1)^ are independently normalized using separate LayerNorm layers. The normalized pair representation is linearly projected to generate a bias term 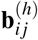 for each attention head *h*. This bias is added to the standard scaled dot-product attention score:

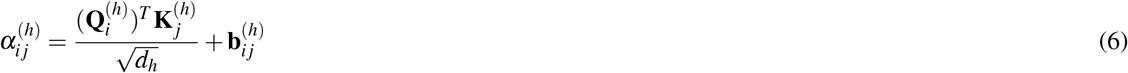

where Query (**Q**), Key (**K**), and Value (**V**) tensors are linear projections of the normalized **E**^(*k*)^. The attention output is aggregated and added back to the input sequence representation via a residual connection to produce **E**^(*k*+1)^:

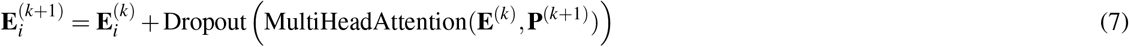

##### (d) Output layer and symmetrization

After *N* blocks, the final pair representation **P**^(*N*)^ is passed through a final LayerNorm and a linear projection head, parameterized by **W**_out_, to produce the logit matrix **Ŷ** ∈ ℝ^*L*×*L*^. To enforce the physical constraint of symmetric base pairing, the final matrix is symmetrized:

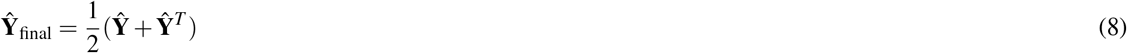

The resulting logits are converted to probabilities ŷ_*i j*_ using a sigmoid function.

### 4.3. Model Training and Loss Function

The ground-truth adjacency matrix *y*, where a **1** indicates the formation of an RNA secondary structure and a **0** indicates its absence, is inherently sparse. To address this significant class imbalance, we trained the model end-to-end using a **weighted binary cross-entropy (BCE)** loss function. A positive weight (*w*) of **300** was introduced to increase the penalty for misclassifying the positive class (i.e., actual RNA structures), giving us the following loss function:

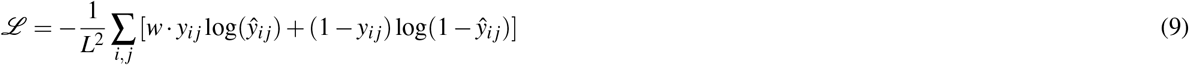

Here, *ŷ*_*i j*_ represents the predicted probability and *y*_*i j*_ is the ground-truth label. The model was trained using the **Adam optimizer** with a learning rate of 1 × 10^−5^. Dropout was applied within the co-evolution blocks to mitigate overfitting, and the entire model was implemented in PyTorch.

### 4.4. Post-processing

After obtaining the symmetric scoring matrix, we follow the post-processing procedures described in e2eFold, BPfold, and UFold^19,20,22^ to generate the final RNA secondary structure prediction. Specifically, the scoring matrix is subjected to the following constraints:

#### Filtering by physical constraints

To enforce structural rules, we allow only canonical (A-U, C-G) and wobble (U-G) base pairs, and prohibit sharp loops (i.e., base pairs between nucleotides less than 4 positions apart). For an RNA sequence *x* = (*x*_1_, *x*_2_, …, *x*_*L*_) of length *L*, we define the set of allowable pairs as:

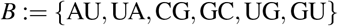

The constraint matrix *P* ∈ {0, 1}^*L*×*L*^ is then defined for all 1 ≤ *i, j* ≤ *L* as:

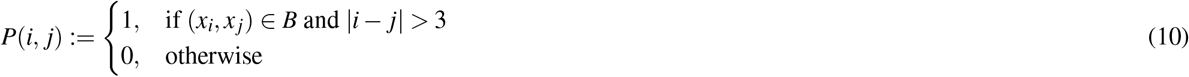

The raw scoring matrix *M* is then symmetrized and filtered:

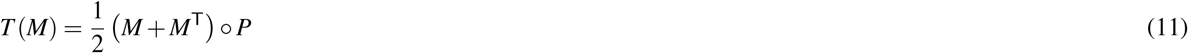

where (·)^T^ denotes matrix transposition and ? is the element-wise product.

#### Ensuring non-overlapping pairs

To ensure that each nucleotide forms at most one base pair, we find the optimal prediction matrix 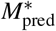 by solving the following constrained optimization problem:

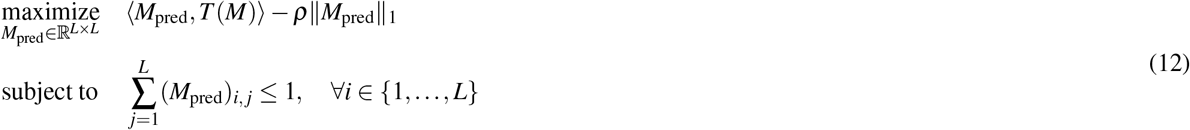

where ⟨··⟩, is the matrix inner product and *ρ* is a regularization coefficient. The resulting matrix 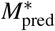 is then thresholded to produce the final binary contact map.

### 4.5. Evaluation Metrics

To evaluate the performance of RNA secondary structure predictions, we adopt three commonly used metrics: Precision, Recall, and F1-score, defined as follows:

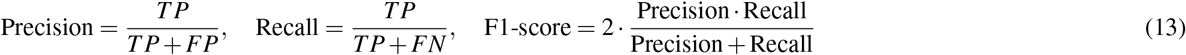

where *TP* (True Positives) denotes the number of correctly predicted base pairs, *FP* (False Positives) denotes the number of incorrectly predicted base pairs, and *FN* (False Negatives) denotes the number of missed base pairs present in the reference structure.

### 4.6. Transfer Learning

We manually collected and carefully curated the PDB-based transfer learning dataset mentioned in SPOT-RNA^18^, resulting in a total of 114 processed RNA sequences (see Section 4.1 for detailed processing procedures). Based on the pretrained model, we conducted transfer learning using this dataset and evaluated the model on our independently constructed test sets: CASP15, CASP16, and PDB-new. Given that the transfer dataset contains only 114 sequences, we adopted a learning rate of `1e-5` and set the loss weight to 50. All other training hyperparameters remained unchanged.

## 5. Data Availability

The datasets used in this study, including TS0, bpRNA-new, CASP15, CASP16, and PDB-new, are publicly available for download from our project website at http://bioinfo.isyslab.info/metafold-rna/download.html.

## 6. Code Availability

The source code for MetaFold-RNA is open-source and available on GitHub at https://github.com/wangleiofficial/MetaFold-RNA. The code is released under an open-source license (see the LICENSE file in the repository for details). The repository includes comprehensive installation guides, usage instructions, and example scripts to reproduce the results presented in the paper. Furthermore, a public web server is available for users to access the core functionalities of MetaFold-RNA without local installation, which can be accessed at http://bioinfo.isyslab.info/metafold-rna/.

## Acknowledgements

This work was supported by National Natural Science Foundation of China under Grant 62172172, Hubei Provincial Natural Science Foundation of China under Grant 2025AFB159, and the Postdoctoral Fellowship Program of CPSF under Grant Number GZC20240545.

## Author contributions statement

L.W. and H.Z. contributed equally to this work and were responsible for the conceptualization, methodology, and initial drafting of the manuscript. Y.W. and Z.X. supervised the project, provided critical revisions, and contributed to data analysis. Z.X. coordinated the research activities, acquired funding, and finalized the manuscript. All authors reviewed and approved the final version of the manuscript.

## Notes

### Competing Interest Statement

The authors have declared no competing interest.

