## Supplementary Material for "MetaFold-RNA: Accurate prediction of RNA secondary structure using a meta-learning-guided deep network"

Table 1: Performance on long-range base pairs (distance $> L/2$) in the TS0 dataset. MetaFold-RNA achieves the highest F1-score (0.741), demonstrating superior accuracy in modeling long-distance structural dependencies.

| **Method** | **Precision** | **Recall** | **F1-score** |
|---|---|---|---|
| LinearFold | 0.736 | 0.641 | 0.545 |
| ContraFold | 0.586 | 0.745 | 0.558 |
| SPOT-RNA | 0.687 | 0.778 | 0.634 |
| UFold | 0.692 | 0.805 | 0.676 |
| EternaFold | 0.593 | 0.744 | 0.563 |
| **MetaFold-RNA** | 0.729 | 0.858 | **0.741** |

Table 2: Performance on long-range base pairs in the bpRNAnew dataset. MetaFold-RNA achieves the highest F1-score (0.753), demonstrating superior accuracy in modeling long-distance structural dependencies.

| **Method** | **Precision** | **Recall** | **F1-score** |
|---|---|---|---|
| LinearFold | 0.850 | 0.755 | 0.677 |
| ContraFold | 0.700 | 0.851 | 0.663 |
| SPOT-RNA | 0.706 | 0.779 | 0.598 |
| UFold | 0.630 | 0.851 | 0.613 |
| EternaFold | 0.731 | 0.844 | 0.661 |
| **MetaFold-RNA** | 0.767 | 0.897 | **0.753** |

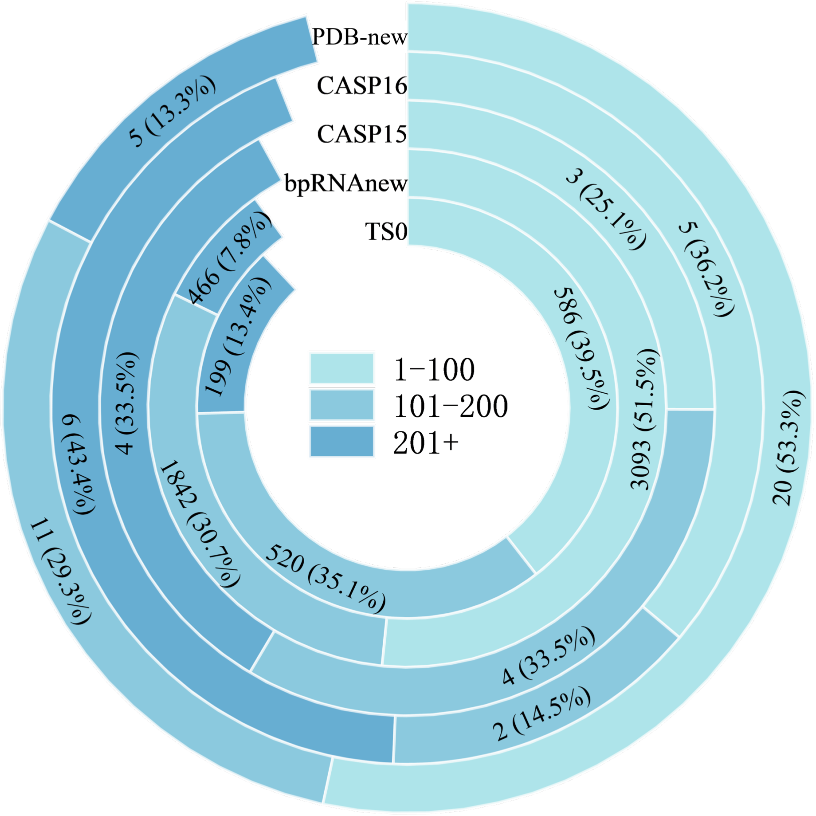


Figure 1: RNA sequence length distribution across the five test datasets (bpRNA-new, TS0PDB-new, CASP15, and CASP16). The stacked bar chart illustrates the number of sequences within three distinct length ranges: 1-100 nt, 101-200 nt, and 201+ nt. The detailed breakdown for each dataset is as follows:

- **bpRNA-new**: This dataset comprises 3093 sequences (51.5%) in the 1-100 nt range, 1842 sequences (30.7%) in the 101-200 nt range, and 466 sequences (7.8%) that are over 200 nt long.
- **TS0**: This dataset contains 586 sequences (39.5%) in the 1-100 nt range, 520 sequences (35.1%) in the 101-200 nt range, and 199 sequences (13.4%) that are over 200 nt long.
- **CASP15**: A total of 11 sequences, with 6 falling into the 1-100 nt range, 2 in the 101-200 nt range, and 3 that are over 200 nt long.
- **CASP16**: A total of 13 sequences, with 5 in the 1-100 nt range, 4 in the 101-200 nt range, and the remaining 4 sequences being over 200 nt long.
- **PDB-new**: This dataset contains 20 sequences (53.3%) in the 1-100 nt range, 11 sequences (29.3%) in the 101-200 nt range, and 5 sequences (13.3%) that are over 200 nt long.
